# Chromosome-level genome assembly of the common chaffinch (Aves: *Fringilla coelebs*): a valuable resource for evolutionary biology

**DOI:** 10.1101/2020.11.30.404061

**Authors:** María Recuerda, Joel Vizueta, Cristian Cuevas-Caballé, Guillermo Blanco, Julio Rozas, Borja Milá

## Abstract

The common chaffinch, *Fringilla coelebs,* is one of the most common, widespread and well-studied passerines in Europe, with a broad distribution encompassing Western Europe and parts of Asia, North Africa and the Macaronesian archipelagos. We present a high-quality genome assembly of the common chaffinch generated using Illumina shotgun sequencing in combination with Chicago and Hi-C libraries. The final genome is a 994.87 Mb chromosome-level assembly, with 98% of the sequence data located in chromosome scaffolds and a N50 statistic of 69.73 Mb. Our genome assembly shows high completeness, with a complete BUSCO score of 93.9% using the avian dataset. Around 7.8 % of the genome contains interspersed repetitive elements. The structural annotation yielded 17,703 genes, 86.5% of which have a functional annotation, including 7,827 complete universal single-copy orthologs out of 8,338 genes represented in the BUSCO avian data set. This new annotated genome assembly will be a valuable resource as a reference for comparative and population genomic analyses of passerine, avian and vertebrate evolution.

## Introduction

The decreasing costs of DNA sequencing, along with advances in computational genomics, are promoting a rapid increase in the availability of high-quality reference genomes of non-model species, which greatly improves our capacity to address a range of biological questions from a genomic perspective. Among them, the correct annotation of protein-coding genes in whole genomes allows to identify new genes involved in the process of evolutionary adaptation and provides a better understanding of the evolutionary mechanisms involved in the speciation process. Avian genomes are particularly suited for studying the molecular basis of speciation as they have a relatively simple architecture and are among the smallest within amniotes, ranging from 0.91 to 1.3 Gb (Gregory 2002). In the last decade, the number of bird reference genomes has increased dramatically (e.g. Dalloul et al. 2010; Warren et al. 2010; Zhang et al. 2012; Jarvis et al. 2014; Poelstra et al. 2014; Frankl-Vilches et al. 2015; Friis et al. 2018; Louha et al. 2019; Peñalba et al. 2019; Ducrest et al. 2020, Wang et al. 2020), providing major scientific breakthroughs in phylogenetics (e.g., Alström et al. 2018; Braun et al. 2019; Jarvis et al. 2015), comparative genomics (Feng et al. 2020), adaptation genomics (Wirthlin et al. 2014; Lawson & Petren 2017), and genomic architecture (Poelstra et al. 2014; Vijay et al. 2016), among others. Moreover, the Ten-Thousand Bird Genomes (B10K) consortium is currently sequencing and assembling over 300 avian genomes that correspond to at least one representative per family (Zhang 2015).

The common chaffinch (Aves, Passeriformes, Fringillidae, *Fringilla coelebs*) is a widely distributed species, ranging from across Eurasia to the north of Africa, and has colonized three Macaronesian archipelagos in the Atlantic Ocean (Azores, Madeira and the Canary Islands) (Collar, Newton & Bonan 2020). With about 15 currently recognized subspecies, the common chaffinch is an ideal system for testing hypotheses on the evolutionary process given its distribution across the continent and the colonization of several oceanic islands, recognized as excellent natural laboratories for studying evolution (Brown et al. 2013). Island systems have inspired the development of biogeographical theories (MacArthur and Wilson 1967) and are of central importance for understanding the role of area and isolation in colonization, extinction and speciation rates (Valente et al. 2020), which are processes influencing global patterns of species richness (Losos and Schluter 2000). Species that have colonized insular environments, like the common chaffinch, are also excellent systems for the study of demographic events, such as bottlenecks leading to small effective population size (Ne) (Leroy et al. 2020), or the roles of drift and selection in the divergence process (Barton 1996). The common chaffinch has been intensively studied using molecular tools, so that the availability of a reference genome represents a valuable resource to improve our understanding of avian evolution, biogeography and demography (Illera et al., 2018).

## Materials and Methods

### Sample collection and genome assembly

A blood sample was extracted from a common chaffinch female captured in Torreiglesias, Segovia, Spain, in 2017 and frozen immediately in liquid nitrogen. The sample was handled by Dovetail Genomics for DNA extraction, sequencing and genome assembly using the HiRise pipeline (Putnam et al. 2016). The absence of a Z chromosome in a first assembly (Genbank assembly accession: GCA_015532645.1) led us to conduct the novel assembly presented here, which includes sex-linked scaffolds used to reconstruct the chaffinch Z chromosome (see File S1 in supplementary materials for specific details). Gene completeness in the chaffinch genome assembly (and in the annotated gene set) was assessed through BUSCO (Benchmarking Universal Single-Copy Orthologs) v4.0.5 (Seppey et al. 2019) by using the 8,338 single-copy orthologous genes in the Aves lineage group odb10, using chicken as the Augustus reference species.

### Identification of repetitive regions and gene annotation

Repetitive regions were identified and masked prior to gene prediction. First, repeats were modelled *ab initio* using Repeat Modeler 1.0.11 (Smit and Hubley 2019) in scaffolds longer than 100 Kb with default options. The repeats obtained were merged with known bird repeat libraries from the RepBase database (RepBase-20181026) (Bao, Kojima, and Kohany 2015), Dfam_Consensus-20181026 and repeats from the zebra finch (obtained from B10K). The resulting repeat library was compared against the complete assembly with Repeat Masker 4.0.7 (Smit, Hubley & Green 2015) and the identified regions were soft-masked. For the identification and description of microsatellites in the common chaffinch genome assembly we used GMATA v.2.01 (Wang & Wang 2016), with sequence motif length between 2 and 20 bp. Gene prediction was conducted with BRAKER v2.1.5 (Hoff et al. 2015) and GeMoMa v1.7.1 (Keliwagen et al. 2016, 2018). First, the conserved orthologous genes from BUSCO Aves_odb10 were used as proteins from short evolutionary distance to train Augustus (Gremme et al. 2005; Stanke et al. 2006; see Figure 3B from Hoff et al. 2019). The predicted proteins were combined with homology-based annotations using the zebra finch (GCF_008822105.2; Warren et al. 2010) and chicken (GCF_000002315.6; Hillier et al. 2014) annotated genes with GeMoMa pipeline, obtaining the final reported gene models. We applied a similarity-based search approach to assist the functional annotation of the chaffinch predicted proteins, using the UniProt SwissProt database, the annotated proteins from the zebra finch genome (Warren et al. 2010; UniProt Consortium 2014) and InterProScan v5.31 (Jones et al. 2014). The functional annotation, including Gene Ontology terms, was integrated from all searches providing a curated set of chaffinch coding genes (see File S1 in supplementary materials for further details).

### Non-coding RNA prediction and identification

For the prediction and functional classification of Transfer RNAs (tRNAs) in the common chaffinch genome we used tRNAscan-SE v2.0 (Lowe & Chan 2016). The tRNA search across the genome and the identification of ncRNA (non-coding RNA) homologues was conducted using the software package Infernal v1.1.1 (Nawrocki 2014) (see File S1 in supplementary materials for details). For comparative purposes, we added our results to those from Louha et al. (2019), which compared different genome assemblies of avian species.

## Results and Discussion

### Assembly and quality control

The total length obtained by the HiRise software for the common chaffinch assembly was 994.87 Mb. Nevertheless, the estimate from *k*-mer metrics is 1.2 Gb. The discrepancy between these estimates could be caused by the presence of repetitive elements given the assembly strategy used, which could have been improved including long-read sequencing technologies. This final assembly consists of 3,255 scaffolds, 3,239 over 1kb and an N50 of 69.73 Mb (Table 1) with a sequence coverage of 249x. The use of Chicago and Hi-C libraries provided a clear improvement in quality by increasing 917 times the scaffold N50, reducing the number of scaffolds from 38,666 to 3,255 (Table 1, see File S1 for details). In fact, 98 % of the total genome sequence maps in the 30 described chromosomes.

**Table 1.** Quality statistics of the different stages of the common chaffinch *(Fringilla coelebs)* assembly.

|  | Shotgun | Chicago HiRise | Dovetail Hi-C |
| --- | --- | --- | --- |
| **Total length** | 903.83 Mb | 907.32 Mb | 994.87 Mb |
| **Scaffold N50** | 0.076 Mb | 8.033 Mb | 69.727 Mb |
| **Scaffold N90** | 0.011Mb | 1.812 Mb | 13.238 Mb |
| **Scaffold L50** | 3,104 | 31 | 5 |
| **Scaffold L90** | 14,974 | 125 | 19 |
| **Longest scaffold** | 1,012,806 bp | 43,054,799 bp | 147,056,713 bp |
| **Number of scaffolds** | 38,666 | 3,767 | 3,255 |
| **Number of scaffolds > 1kb** | 38,666 | 3,767 | 3,255 |
| **Contig N50** | 67.15 kb | 67.08 kb | 66.81 kb |
| **Number of gaps** | 8,240 | 43,147 | 49,330 |
| **Percent of genome in gaps** | 0.11 % | 0.50 % | 0.49 % |

The chaffinch genome showed high synteny with the zebra finch genome (Fig. 1), evidencing the completeness of the assembly, with all micro-chromosomes and the Z chromosome present in the assembly. In addition, the alignment between these genomes suggests the presence of several inversions in chromosomes 1, 1A, 2, 3, 5, 7, 8 and 9. Several studies have documented that inversions are very common in birds (Aslam et al. 2010, Völker et al. 2010, Skinner and Griffin 2011, Zang et al. 2014). For instance, Hooper and Price (2017) identified 319 inversions on the 9 largest autosomes combined in 81 independent clades. No putative contaminations were detected and 89.6% of the reads were mapped in the genome assembly (Fig. S1). The mean GC content of the assembly was 41.86% (±11 SD). The common chaffinch genome assembly included 7,832 complete copies (93.9%) out of the 8,338 BUSCO dataset from avian genomes, among which 7,816 were single-copy orthologs and 16 were duplicated. Only 1.8% of the gene models were fragmented, and 4.3% were missing in the genome. These few missing gene models could represent divergent or lost genes in our species, but also could be related with putative errors during the assembly process or missing data.

**Figure 1.**
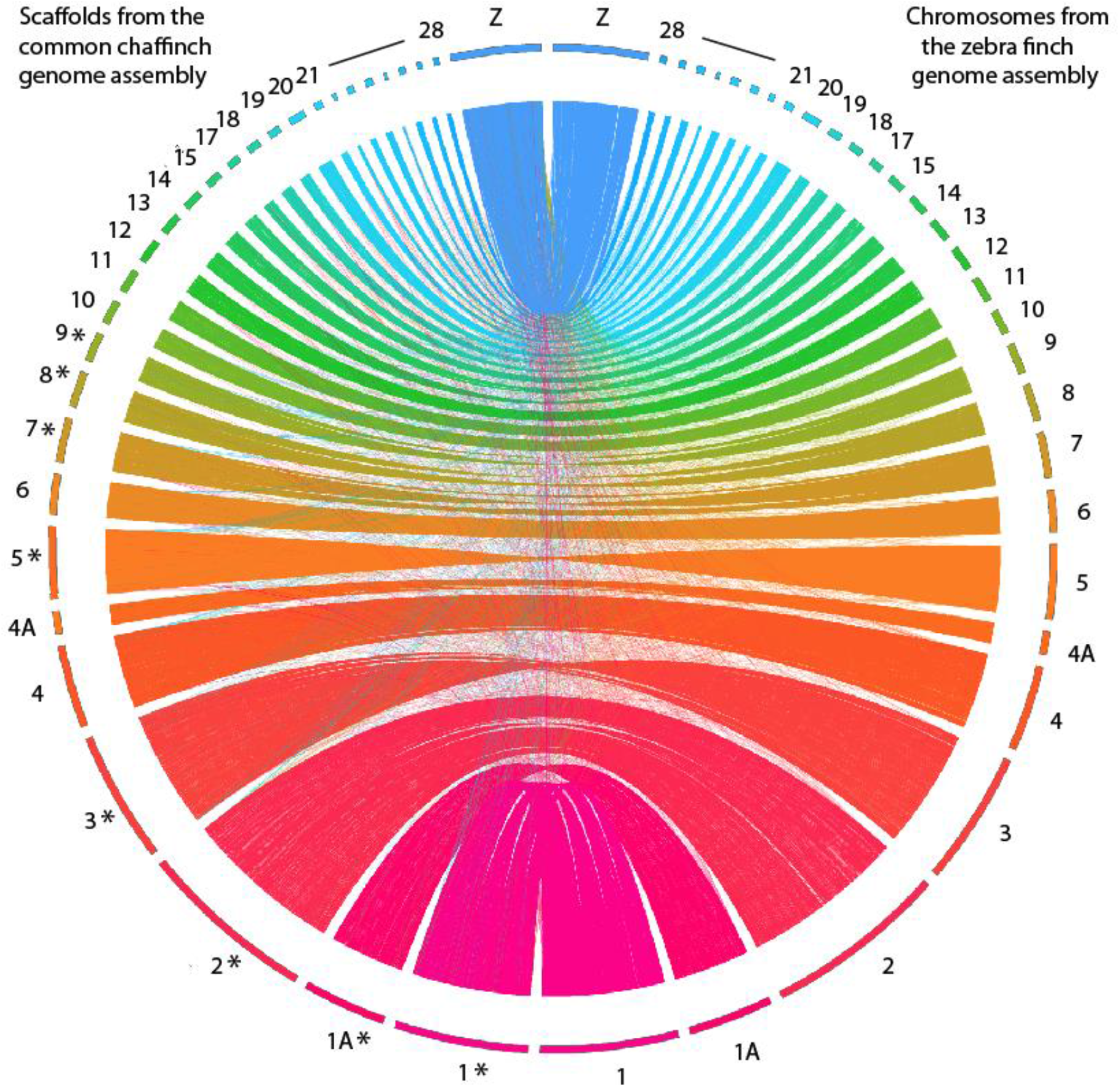
Circos plot comparing the zebra finch (right) and the common chaffinch (left) genome assemblies (Krzywinski et al. 2009). The alignment was conducted using nucmer (Marçais et al. 2018). The common chaffinch chromosomes marked with an asterisk (*) show inversions with respect to the zebra finch assembly.

### Repetitive regions

Overall, 7.82% of the genome assembly are repeats (~78 Mb), of which 85.4 % are transposable elements (TEs). The most abundant TEs are LINEs (53.5%) followed by LTR (29.4%), DNA elements (4.1 %) and SINEs (1.4 %), with the remaining 11.6% unclassified. The rest of repeats (14.6 %) contained simple repeats (75.4%), low complexity repeats (18.5 %), satellites (4.2%) and small RNA (1.9 %) (Table S1). The number of repetitive regions is within the expected range in birds, which is at 4 to 10% of the genome (Zhang et al. 2014).

A total of 111,076 microsatellites, with motif length ranging between 2 and 20 bp, were identified in the common chaffinch genome (File S2: microsatellites and their genomic locations). The most common *k*-mer sizes conforming the microsatellites were 2 (68.2%), 3 (15.9%) and 4 (8.2%) (File S2). The most common length of the microsatellites was 10 bp (40.4%), followed by 12 bp (13%) and 15 bp (8.8%) (see File S2 for the length distribution of microsatellites). In addition, the number of microsatellites was positively correlated with the sequence length (File S2; also included the frequency of occurrence in every scaffold).

### Gene annotation and function prediction

Our annotation pipeline combining both *de novo* and homology-based predictions inferred 21,831 proteins encoded by 17,703 genes in the common chaffinch genome with a mean length of 15,818 bp (Table 2). The common chaffinch genome annotation (File S3, annotation file in general feature format) included 7,850 complete copies (94.2%) out of the 8,338 of BUSCO avian dataset used, retrieving all expected copies with a slight increase from that estimated in the un-annotated genome (see above). Among the complete BUSCO genes, 7,827 were single-copy orthologs (99.7%) and 23 were duplicated (0.3%). Around 1.9% (162) of the gene models were fragmented and 3.9% showed no significant matches (326).

**Table 2.** Genome statistics and predicted ncRNAs of the *F. coelebs* genome compared to other similarly sized avian species *(Melospiza melodia, Taeniopygia guttata, Ficedula albicollis, Manacus vitellinus,* and *Geospiza fortis),* modified from Louha et al. (2019).

|  | *F. coelebs* | *M. melodia* | *T. guttata* | *F. albicollis* | *M. vitellinus* | *G. fortis* |
| --- | --- | --- | --- | --- | --- | --- |
| Number of genes | 17,703 | 15,086 | 17,561 | 16,763 | 18,976 | 14,399 |
| Mean gene length (bp) | 15,818 | 14,457 | 26,458 | 31,394 | 27,847 | 30,164 |
| Number of CDSs | 17,703 | 15,086 | 17,561 | 16,763 | 18,976 | 14,399 |
| Mean CDs length (bp) | 1,679 | 1,325 | 1,677 | 1,942 | 1,929 | 1,766 |
| Number of exons | 221,872 | 131,940 | 171,767 | 189,043 | 190,390 | 164,721 |
| Mean exon length (bp) | 165 | 153 | 255 | 253 | 264 | 195 |
| Mean number of exons/gene | 10.16 | 8.67 | 10.25 | 12.22 | 11.51 | 11.41 |
| Number of introns | 200,041 | 116,724 | 153,909 | 171,236 | 171,089 | 149,563 |
| Mean intron length (bp) | 1,902 | 1,695 | 2,930 | 3,257 | 3,294 | 2,813 |
| Total proteins | 21,831 |  |  |  |  |  |
| **ncRNA** |  |  |  |  |  |  |
| tRNA | 325 | 267 | 184 | 179 |  |  |
| miRNA | 140 | 166 | 302 | 510 |  |  |
| snRNA | 18 | 16 | 44 | 32 |  |  |
| snoRNA | 126 | 154 | 241 | 199 |  |  |
| rRNA | 5 | 8 | 100 | 22 |  |  |
| lncRNA | 17 | 20 | 908 | 1473 |  |  |

Over all predicted proteins, 19,458 (89.1%) provided positive BLASTP hits against the Uniprot SwissProt database, and 19,617 (89.9%) against the annotated proteins from the zebra finch genome. In addition, InterproScan identified 18,551 (85%) specific protein-domain signatures in the predicted peptides. The combination of the annotation from these databases allowed assigning a functional annotation with GO terms to 19,425 proteins (89%) assigned to 15,309 genes (86.5%; File S4).

### tRNAs and other non-coding RNA prediction

The search by tRNAscan-SE (File S5) identified 325 tRNAs in the common chaffinch genome, of which 167 decode for the standard twenty amino acids. Among all the tRNAs detected, 131 presented low scores and therefore were categorized as pseudogenes (i.e. lacking tRNA-like secondary structures). There were no suppressor tRNAs, 1 had undetermined isotopes, 25 were chimeric and 15 included introns within their sequences. One of the tRNAs was predicted to code for selenocysteine (sequences and structures of the predicted tRNAS are available in File S6). In addition, the search against both tRNA databases (GtRNAdb and tRNAdb) yielded positive results in many other species, suggesting that tRNA prediction in our assembly was correct. Moreover, our searches using Infernal identified 354 ncRNAs, which were classified as follows: 39 CREs, 2 Ribozymes, 7 Gene, 140 miRNAs, 126 snoRNAs, 18 snRNAs, 5 rRNAs, and 17 lncRNAs (File S7). The number of tRNAs predicted in the common chaffinch genome is the highest when compared to other passerine species (i.e. *M. melodia, T. guttata* and *F. albicollis),* but the other types of ncRNAs present similar values to the *M. melodia* genome and lower than the other two species (Table 2), probably because we applied a strict threshold to avoid an excess of false positives.

## Conclusions

We provide here a high-quality assembly for the common chaffinch, a valuable resource as a reference genome to address a range of biological questions from a genomic perspective. Moreover, our annotation provides useful information to detect candidate genes involved in adaptation and divergence processes. The combination of the Chicago and shotgun sequencing with the HiRise assembly approach lead to a highly contiguous chromosome-level genome assembly. The genome assembly size was 994.87 Mb, with the 30 chromosomes accounting for 98% of it. Although the expected length of the genome was 1.2 Gb, closer to those obtained in other avian species by flow cytometry (Gregory 2002), the BUSCO analyses showed that both the assembly and structural annotation encode 93.9% and 94.2% complete copies out of the 8,338 orthologous conserved genes in avian species, respectively. This discrepancy of the genome size could be caused by the absence of repetitive elements in the assembly. The structural annotation predicted a high number of genes compared to other avian genomes, with a high percentage of the genes (86.5%) assigned to functional annotation and GO terms.

## Acknowledgements

The research was supported by grant CGL2015-66381P from the Spanish Ministry of Economy and Competitiveness to BM and GB, and grant PGC2018-098897-B-I00 from the Spanish Ministry of Science and Innovation to BM. MR was supported by a doctoral fellowship from the Spanish Ministry of Education, Culture, and Sport (FPU16/05724).

## Data availability

The chaffinch genome assembly and the raw data have been deposited at NCBI under BioProject PRJNA674347 with accession number JADKPM000000000, and all described datasets including the annotation are publicly accessible in *Figshare* (https://doi.org/10.6084/m9.figshare.13296122.v1).

